# Comparative phylogenomic analysis reveals evolutionary genomic changes and novel toxin families in endophytic *Liberibacter* pathogens

**DOI:** 10.1101/2021.06.02.446850

**Authors:** Yongjun Tan, Cindy Wang, Theresa Schneider, Huan Li, Robson Francisco de Souza, Xueming Tang, Tzung-Fu Hsieh, Xu Wang, Xu Li, Dapeng Zhang

## Abstract

*Liberibacter* pathogens are the causative agents of several severe crop diseases worldwide, including citrus Huanglongbing and potato Zebra Chip. These bacteria are endophytic and non-culturable, which makes experimental approaches challenging and highlights the need for bioinformatic analysis in advancing our understanding about *Liberibacter* pathogenesis. Here, we performed an in-depth comparative phylogenomic analysis of the *Liberibacter* pathogens and their free-living, nonpathogenic, ancestral species, aiming to identify the major genomic changes and determinants associated with their evolutionary transitions in living habitats and pathogenicity. We found that prophage loci represent the most variable regions among *Liberibacter* genomes. Using gene neighborhood analysis and phylogenetic classification, we systematically recovered, annotated, and classified all prophage loci into four types, including one previously unrecognized group. We showed that these prophages originated through independent gene transfers at different evolutionary stages of *Liberibacter* and only the SC-type prophage was associated with the emergence of the pathogens. Using ortholog clustering, we vigorously identified two additional sets of genomic genes, which were either lost or gained in the ancestor of the pathogens. Consistent with the habitat change, the lost genes were enriched for biosynthesis of cellular building blocks. Importantly, among the gained genes, we uncovered several previously unrecognized toxins, including a novel class of polymorphic toxins, a YdjM phospholipase toxin, and a secreted EEP protein. Our results substantially extend the knowledge on the evolutionary events and potential determinants leading to the emergence of endophytic, pathogenic *Liberibacter* species and will facilitate the design of functional experiments and the development of new detection and blockage methods of these pathogens.

**Importance:** *Liberibacter* pathogens are associated with several severe crop diseases, including citrus Huanglongbing, the most destructive disease to the citrus industry. Currently, no effective cure or treatments are available, and no resistant citrus variety has been found. The fact that these obligate endophytic pathogens are not culturable has made it extremely challenging to experimentally uncover from the whole genome the genes/proteins important to *Liberibacter* pathogenesis. Further, earlier bioinformatics studies failed to identify the key genomic determinants, such as toxins and effector proteins, that underlie the pathogenicity of the bacteria. In this study, an in-depth comparative genomic analysis of *Liberibacter* pathogens together with their ancestral non-pathogenic species identified the prophage loci and several novel toxins that are evolutionarily associated with the emergence of the pathogens. These results shed new lights on the disease mechanism of *Liberibacter* pathogens and will facilitate the development of new detection and blockage methods targeting the toxins.

## Introduction

*Candidatus* Liberibacter, a genus of Gram-negative bacteria in the order of Rhizobiales, has recently received increasing attention due to the fact that several of its species are associated with severe diseases on multiple crop plants, such as citrus, potato, and tomato. These include three species, *Candidatus* Liberibacter asiaticus found in Asia and North America, *Candidatus* Liberibacter africanus found in Africa ^1^, and *Candidatus* Liberibacter americanus found in Brazil ^2^, that cause the citrus Huanglongbing (HLB) disease, and *Candidatus* Liberibacter solanacearum, that causes similar diseases in tomato and potato, referred to as Psyllid yellows and Zebra Chip (ZC), respectively ^3,4^ The HLB disease, also known as citrus greening, is the most destructive, worldwide disease of citrus ^5^. It is characterized by yellowing of citrus tree leaves, premature defoliation, decay of feeder rootlets and lateral roots, production of small bitter fruit, and eventually death of the citrus tree ^6^. Thus, HLB has led to a substantial loss of citrus production and severe damage to the economy and job market ^7^. Unfortunately, there is currently no effective cure or treatments available, and no resistant citrus variety has been found so far. In a similar vein, ZC is a new disease of potato that was first identified in Mexico in 1994 ^8^ and has quickly spread to many countries in recent years. It negatively affects growth, yield, propagation potential, and qualities of tubers and thus impacts international trade and the economy significantly ^9^.

Given the severity and rapid spread of these diseases in commercial crops, extensive research efforts have been made to better understand the basic biology of these pathogens and the pathogenesis mechanisms of the diseases, especially on HLB. However, the progress has been slow due to two main obstacles. First, the *Liberibacter* pathogens of these diseases are endophytic bacteria, whose transmissions are naturally vectored by several psyllids, such as Asian citrus psyllid (*Diaphorina citri*)^10^, African citrus psyllid (*Trioza erytreae*)^11^, and *Bactericera cockerelli* ^9^. Therefore, they are not culturable which makes it difficult for wet-lab researchers to conduct functional studies. Second, secretion of toxins or effectors represents one of the most important mechanisms of bacterial pathogenesis and virulence. These toxins either manipulate host/vector immunity and physiology or damage the host cells ^12^. However, such molecules involved in the HLB disease have remained elusive for years. Previous research has focused primarily on secreted proteins in HLB-associated pathogens which carry T2SS specific signal peptides ^13^. Thus far, the only proteins claimed to be the potential toxins behind the HLB disease have been the short proteins carrying these signal peptides ^14–17^. However, these proteins are mainly present in one strain of the *C*.L. asiaticus species, and not found in other *C*.L. asiaticus strains and other *Liberibacter* pathogens associated with HLB (*C*.L. americanus and *C*.L. africanus)^14,16–21^, suggesting that other types of toxins or effectors may contribute to the primary HLB pathology. Without accurate and comprehensive information about the HLB-associated toxins/effectors, targeted strategies for detection, prevention, and more importantly blockage will not be possible.

In this study, we aimed to tackle these problems by utilizing the available genome information and dedicated bioinformatics strategies. Our analysis is based on the observed phenotypic difference between the most ancestral *Liberibacter* species, *Liberibacter crescens*, that displays a free-living habitat and is apparently non-pathogenic, and the descendants that are both endophytic and pathogenic. We hypothesized that the functional difference is determined by the genomic changes, including both gene-loss and gene-gain events, that occurred at the common ancestor of the pathogens, after its divergence from *L. crescens*. We have designed multiple comparative genomics strategies to systematically mine the major genomic differences, including unique prophage loci and other genes that are evolutionarily associated with the emergence of the pathogens. More importantly, we identified several novel toxin proteins, including a new type of polymorphic toxins, a YdjM phospholipase toxin, and a secreted EEP family protein. The new information gained from this research provides important insights into the evolution and pathogenesis of *Liberibacter* pathogens and will facilitate future research to develop novel detection and blockage methods targeting the toxins.

## Results

### Phylogenetic relationships of HLB-associated pathogens suggest an evolutionary transition from the non-pathogenic ancestor to pathogenic descendants

We first sought to understand the evolutionary relationship of the three HLB-associated pathogens. We collected all *Liberibacter* species whose genome information is available in the NCBI GenBank database (Fig. 1a and Supplementary Table 1). This includes eight genomes of HLB-associated pathogens (six strains of *C*.L. asiaticus ^5,22–26^, one of *C*.L. africanus ^27^, and one of *C*.L. americanus ^2^), one genome of *C*.L. solanacearum^28^, one genome of *C*.L. europaeus, which is a potential pathogen of *Cytisus scoparius* and vectored by *Arytainilla spartiophila* ^29^, and two genomes of *L. crescens*, which were isolated from papaya and represent the most basal lineage within the Liberibacter genus^30,31^. We conducted Maximum Likelihood phylogenetic analyses by using three conserved genes including 16S rRNA, 23S rRNA, and DNA polymerases A, all of which support the same tree topology (Fig. 1b-d). Importantly, this analysis revealed that the species associated with HLB are not monophyletic. Both *C*.L. africanus and *C*.L. asiaticus share a common ancestor. However, *C*.L. americanus shows a close relationship with *C*.L. europaeus. Between these two groups is *C*.L. solanacearum, the pathogen of potatoes, tomatoes, and carrots. Further, all five species of pathogens share a common ancestor, whose sister group is *L. crescens*. *L. crescens* are free-living bacteria and do not seem to be pathogenic, despite the fact that they were isolated from papaya ^31,32^. Our phylogenetic trees also support its ancestral (deepest branching) position at the base of *Liberibacter* (Fig. 1b-d).

**Fig. 1.**
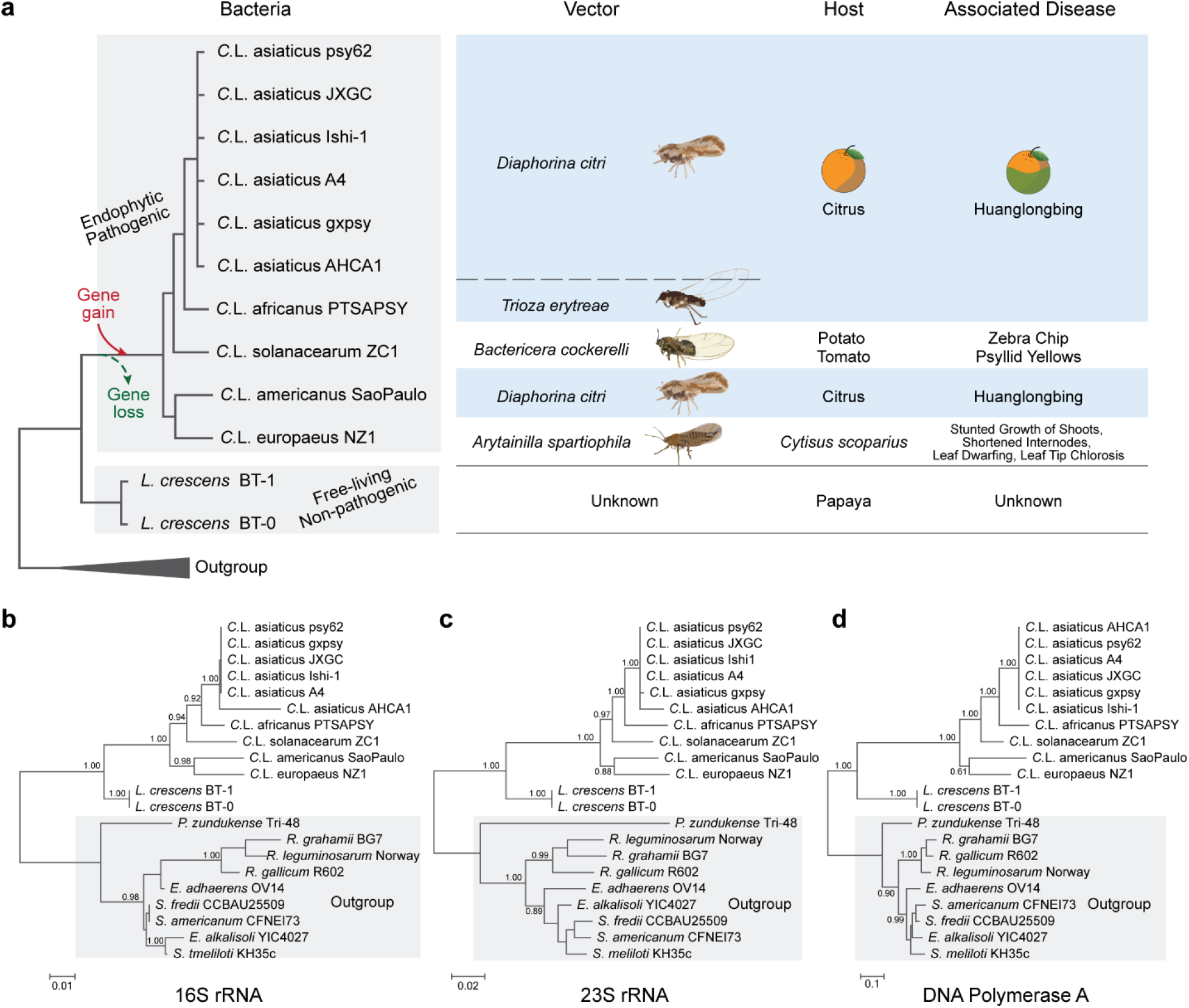
Phylogenetic relationship of HLB-associated bacteria and their relative species. **a** Detailed information of *Liberibacter* species whose genomes were investigated in this study, including their evolutionary relationship, transmitted vectors, hosts, and associated diseases. The evolutionary relationship was derived from phylogenetic trees of 16S rRNA (**b**), 23S rRNA (**c**), and DNA polymerase A (**d**). **b-d** The phylogenetic trees were inferred using the Maximum Likelihood (ML) method, where the supporting values from 100 bootstrap are shown for major branches only. The outgroup clades are colored in grey background.

According to this phyletic pattern between the ancestral species of *Liberibacter*, *L. crescens*, which is free-living and apparently non-pathogenic, and the descendants, which are endophytic and pathogenic bacteria, there is a clear transition in term of living habitats and pathogenicity. In other words, both the intracellular habitats and pathogenicity of these bacteria are derived traits of *Liberibacter* during evolution. We hypothesize that the development of these new traits was attributed to the genomic changes, both gene-loss and gene-gain events, that occurred in the ancestor of the *Liberibacter* pathogens. With this hypothesis, we designed several phylogenomic comparison strategies to specifically identify the genes or genomic regions that display unique phyletic patterns in either pathogenic descendants or ancestral *L. crescens* species and to understand the evolution and pathogenicity mechanisms of *Liberibacter*.

### Whole genome comparisons of *Liberibacter* bacteria

The first computational strategy involved comparing whole *Liberibacter* genomes in order to develop a general idea of genomic dynamics and to identify any large genomic regions that were deleted or inserted during evolution. Pairwise TBLASTX comparisons were used to identify the gene correspondence between the genomes (Fig. 2). Each line indicates one correspondence of the homologs. We found that between five pathogenic species, the majority of their genome regions preserve similar gene composition and organization. The major difference between their genomes arises from large-scale genome recombination and inversions (as shown in crossed blue lines). However, when we compared genomes of the non-pathogenic *L. crescens* and pathogenic *C*.L. europaeus, the difference between their genomes is evident. Although many short operonic structures are still preserved between the two genomes, the recombination and inversions are more prominent and extensive. Thus, the observed extensive genomic changes between the non-pathogenic ancestor and pathogenic ancestor are consistent with their transitions in living habitat and pathogenicity.

**Fig. 2.**
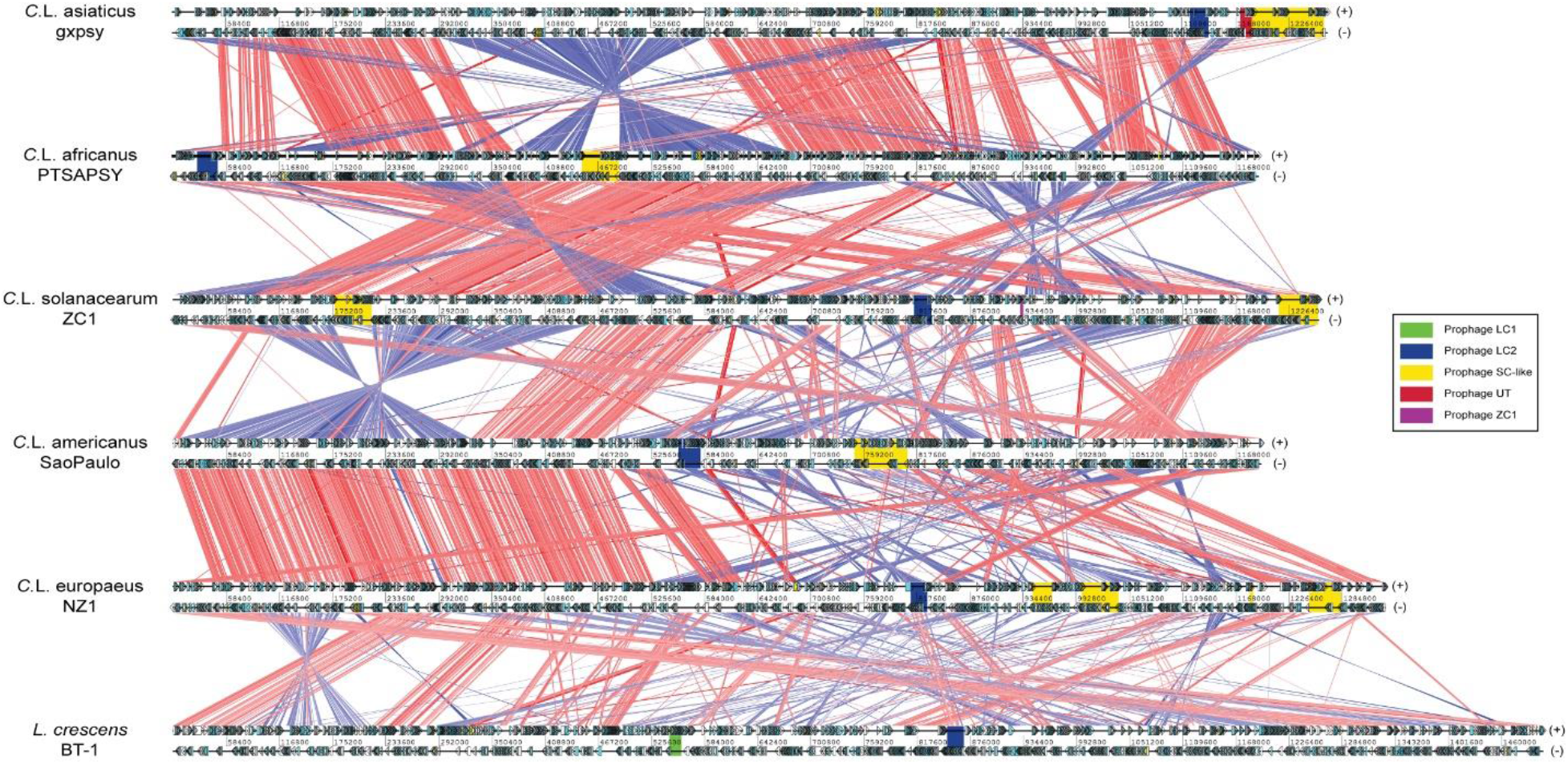
Whole genome comparisons of *Liberibacter* species. The genomes are labelled by their scientific names followed by strain information. Cyan and yellow boxes are the protein coding frames and RNA regions on either forward (+, top) and reverse (-, bottom) strands, respectively. Homologous genes between genomes are linked by lines, where the red and blue lines represent the forward and reverse (complementary) matches, respectively. The intensity of the color bands is proportional to the percent identity of the match, where higher intensity indicates higher sequence identity. The phage loci are shown in colored boxes (legend on the right).

When the genome regions that display strong variations among pathogens and between pathogens and the non-pathogenic ancestor were examined, we found that many of them are prophage loci (as highlighted on each genome) (Fig. 2). This is very interesting, as prophages are known to play a critical role in bacterial pathogenesis ^33^ and many toxins or virulence factors of bacterial pathogens are carried by prophage loci, such as cholera toxins ^34^ and the toxins used by pathogenic *E.coli* ^35^. Therefore, we conducted a detailed analysis of all prophage loci to test if any prophage loci are associated with the pathogenicity of the pathogens.

### Genomic organization dissection and classification of *Liberibacter* prophage loci

Several prophage loci have been previously identified in *Liberibacter* bacteria ^30,36,37^. However, their evolutionary relationships, origins, genomic organizations, and functional compositions remain unclear. Thus, we conducted an in-depth gene-neighborhood analysis to systematically extract the potential prophage loci, identify their boundaries, and annotate their components. As a result, we retrieved 36 prophage loci from the 12 available *Liberibacter* genomes (Fig. 3, Supplementary Table 2 and Data 1). We annotated their gene components by clustering the protein products and dissecting their domains. We found that these prophage loci display a highly variable gene composition. However, based on the shared gene components, we were able to classify them into four major types, including LC1, LC2, SC, and UT (Fig. 3). Specifically, the LC1 type is only found in non-pathogenic *L. crescens* whereas LC2 is present in all *Liberibacter* species. They are similar in terms of gene composition; however, LC2 contains several unique genes such as LC_TM, Peptidase_S74_2, LC_3, HTH_XRE, and DUF1376. The SC type is found in all *Liberibacter* pathogens; it typically has a large genome size with big variations among loci which are mainly caused by independent genome deletion and insertions. It is noteworthy that several genomes contain multiple copies of the SC-type phages. These include two copies in *C*. L. asiaticus gxpsy genome, two copies in *C*. L. solanacearum ZC1 genome, and three copies in *C.* L. europaeus NZ1 genome. By contrast, the UT type represents a novel group of prophages that we recovered in all *C*. L. asiaticus strains. Its size is much smaller, and according to its gene composition, it is close to the SC type. There is also a small prophage locus ZC1 on the *C*.L. solanacearum genome, but it is more likely a fragment.

**Fig. 3.**
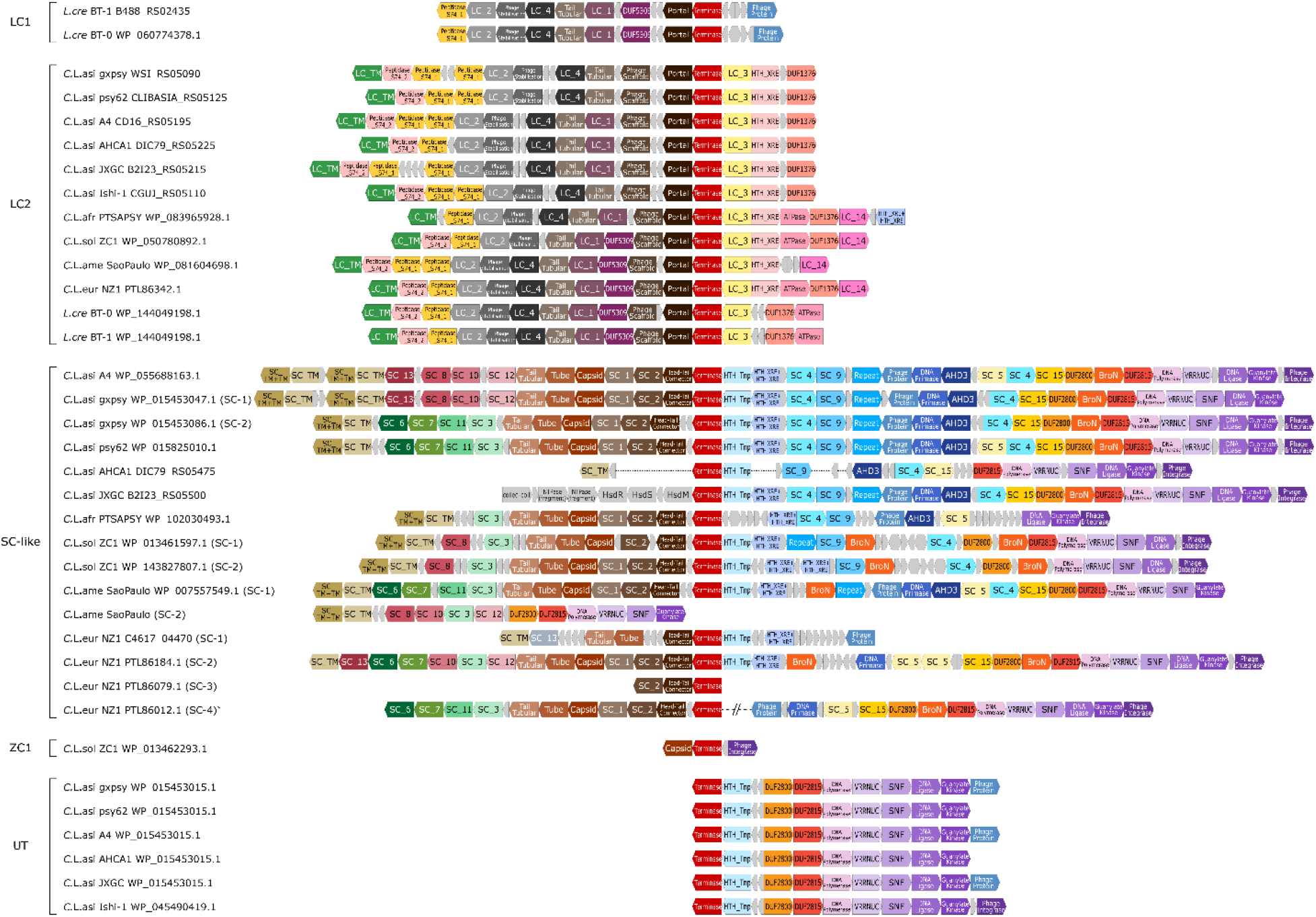
Genomic structures of the prophages identified in the *Liberibacter* species. The coding frames in prophage loci are presented in blocks. The blocks, labelled with gene annotations and highlighted in different colors, are the coding products shared by at least four prophage loci, while the small grey blocks represent non-conserved coding products or pseudogenes. The genome structures were aligned based on the shared terminases of these prophage loci. On the left side, the prophage loci are indicated by their species names, strains, and terminase accession numbers or locus tag. The prophage loci were classified into four types (LC1, LC2, SC-like, UT) and a distinct prophage fragment found in a *C*.L. solanacearum ZC1 strain. One *C*.L. europaeus SC-like prophage structure was derived from two genome contigs, PSQJ01000015.1 and PSQJ01000003.1, which is indicated by a star (*).

### Phylogenetic analysis reveals independent gene transfers of phages to *Liberibacter*

We next attempted to trace the evolutionary histories of these prophages to examine their correlations with the development of pathogenicity. By comparing gene components of prophage loci, we found that terminase, the key phage component involved in the phage DNA packing process ^38^, is the only gene family conserved across all identified prophage loci (Fig. 3). Therefore, we used the terminase as a marker to study the evolutionary origins of these prophages. We conducted several BLASTP searches using the terminases from *Liberibacter* species to collect homologs in the NCBI NR database. Using both Maximum Likelihood and Bayesian Inference analyses (Fig. 4), we found that the terminases of *Liberibacter* prophages are not monophyletic; instead, terminases of different types, LC1, LC2, and SC (together with UT), were nested in three separate clades. This indicates that the LC1, LC2, and SC prophages have different evolutionary origins, and they have been transferred independently to *Liberibacter* at different evolutionary time points (Fig. 4a). In addition to the independent origins, the tree also suggests that multiple copies of the SC phage in several *Liberibacter* pathogens were likely generated from genome-specific duplications, such as those in *C*.L. solanacearum, *C*.L. europaeus, and *C*.L. asiaticus (Fig. 4a). However, the SC prophages also appear to have undergone recombination between species and strains, given the facts: 1) their terminases did not follow a typical pattern of vertical evolution, unlike the LC2 terminases (Fig. 4a); 2) in our genome comparison analysis, the SC phages from different *C*.L. asiaticus strains display a strikingly rapid divergence in certain regions, in contrast to other genomic regions (Fig. 4b).

**Fig. 4.**
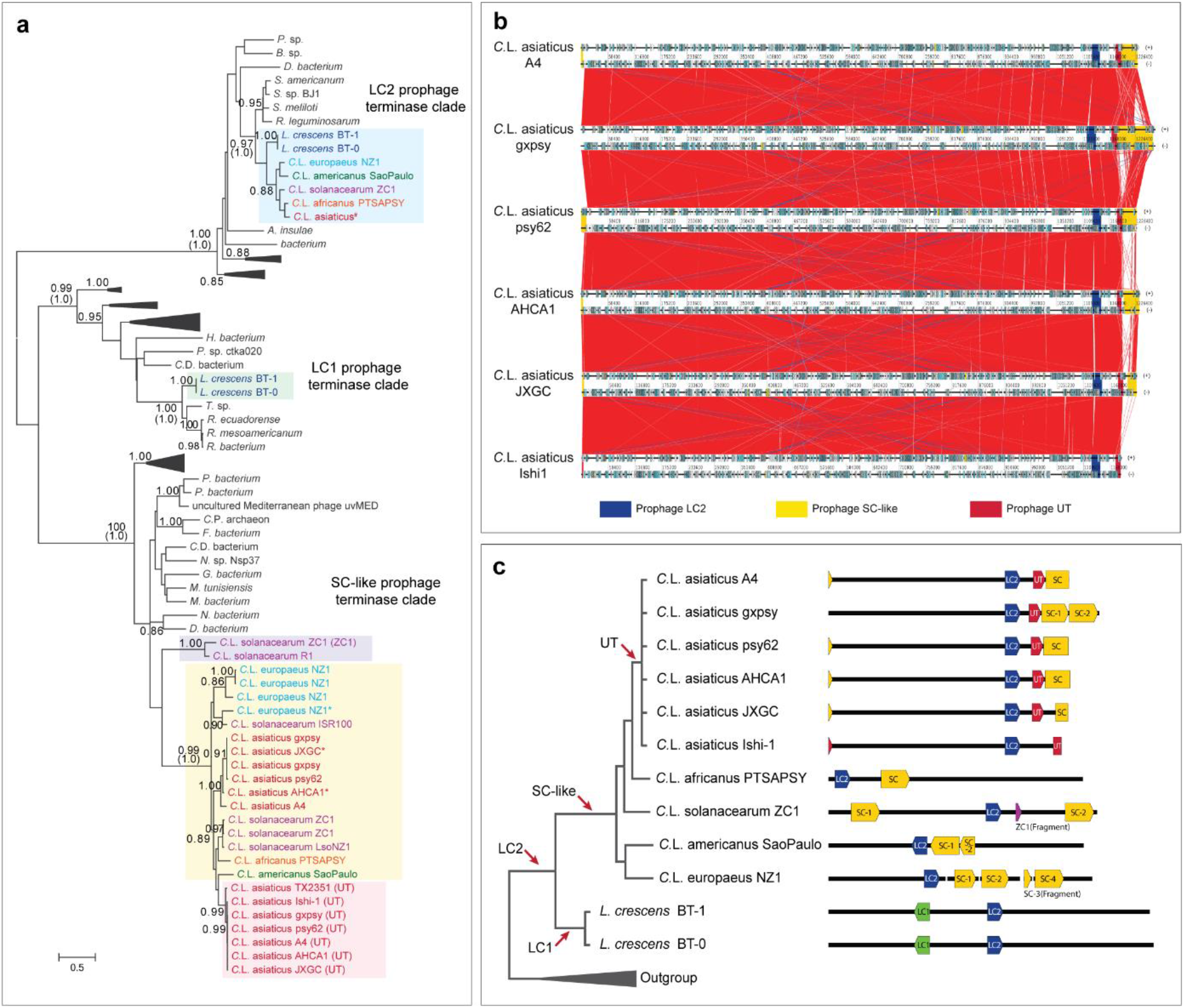
Independent origins of *Liberibacter* prophages. **a** Phylogenetic relationship of prophage terminases. The phylogeny was constructed by using Maximum Likelihood (ML) and Bayesian Inference (BI) methods. The ML tree with the highest log likelihood is presented with the major branches supported by both ML bootstrap values (top) and the BI Posterior values (bottom, bracketed). The terminases from the previously-defined *Liberibacter* prophage types are highlighted in different background color: LC2 prophage in light blue, LC1 prophage in light green, ZC1 prophage in light purple, UT prophage in light red, and SC prophage in yellow. Sequence annotations were colored differently according to their species. The terminase sequences from *C*. L. asiaticus strains are identical, therefore, one sequence is used to represent and indicated by hashtag (#). Several protein sequences were translated by the ORFfinder program, which are indicated by a star symbol (*). **b** Whole genome comparison of different strains of *C*. L. asiaticus. Homologous genes between genomes are linked by lines. Colored boxes are the prophage loci and the high variation of SC-type prophage loci is shown at the 3’ end of the genomes. **c** The inferred independent origins of different *Liberibacter* phage types. The evolutionary relationship of *Liberibacter* species is shown on the left and their composition of the phage loci are illustrated on the right. The coloring themes refer to the Figure 2 legend. The red arrows indicate the potential gene transfer events of different prophage types.

Based on the results from both gene neighborhood-based classification and phylogenetic analysis, it is likely that LC2, LC1, and SC prophages were introduced at the base of *Liberibacter*, the base of non-pathogenic *L. crescens*, and the common ancestor of the *Liberibacter* pathogens, respectively. As for the UT phage, it likely originated from a duplication event of the ancestral SC phage at the base of the *C*.L. asiaticus species (Fig. 4b). According to the phylogenetic histories of these prophages, we can infer that the origin of the SC type of prophages was associated with the emergence of pathogenicity of the *Liberibacter* pathogens.

### Ortholog clustering analysis reveals additional unique genes which were either lost or gained in the ancestor of pathogens

In addition to prophages, we sought to identify other genomic (non-prophage) genes which might be associated with the transition from the non-pathogenic ancestor to pathogenic descendants. These genes should be featured by having undergone either gene-loss or gene-gain events in the common ancestor of all pathogens. Therefore, we were specifically interested in identifying two groups of genes which display unique phyletic patterns: first, the genes, that underwent gene loss, are present in the non-pathogenic *L. crescens* species, but not in pathogenic descendants; second, the genes, that underwent gene gain, are present in all pathogens but not in non-pathogenic *L. crescens* species (Fig. 5a).

**Fig. 5.**
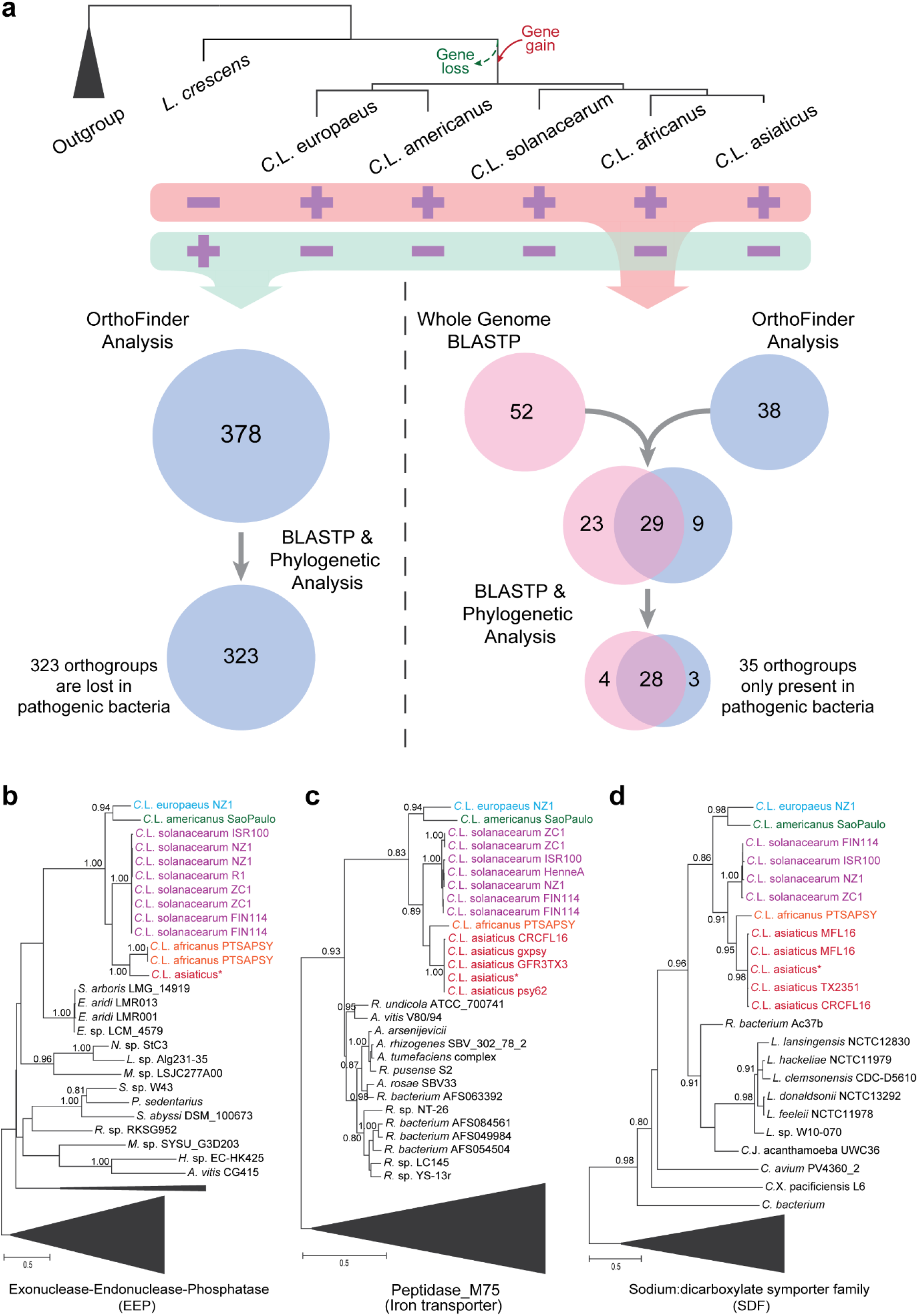
Ortholog clustering analysis identified the genomic genes that were gained and lost during the transition from non-pathogenic ancestor to pathogenic decedents. **a** A conceptual figure illustrating the strategies to identify the *Liberibacter* genes which were gained or lost during evolution according to the phyletic patterns (presence or absence) of orthologous genes in both ancestral free-living *L. crescens* and pathogenic decedents. **b-d** Phylogenetic verification of gene transfer events of three gene products, exonuclease-endonuclease-phosphatase (EEP), peptidase M75 (iron transporter), and sodium:dicarboxylate symporter family (SDF) protein, to the pathogenic ancestor. The trees were inferred using Maximum Likelihood (ML) method, and the supporting values from 100 bootstraps are shown for major branches. Sequences were annotated by their species and strains, colored accordingly. Some sequences from different *C*. L. asiaticus stains are identical and so we use one sequence to represent, which is indicated by a star symbol (*).

To identify these genes, we utilized three steps of analysis. First, we carried out an ortholog analysis using the OrthoFinder program ^39^ to cluster the proteins (excluding the prophage components) of all collected *Liberibacter* genomes into different orthologous groups (orthogroups). Based on their phyletic profiles, we identified the orthogroups that are only present in pathogens (but not in the non-pathogenic ancestor) or in the non-pathogenic ancestor (but not in pathogens). We also utilized whole-genome BLASTP searches to specifically identify the cases of gene gain or gene loss by comparing the *Liberibacter* proteins with other sequences in the NCBI database, according to the pairwise sequence similarity scores. Finally, to validate and confirm the above results, we conducted extensive phylogenetic analyses on all identified orthogroups. From these analyses, we identified 323 orthogroups (about 335 genes in *L. crescens* BT-1) that were lost in all pathogenic descendants and 35 orthogroups (about 35-37 genes in each pathogen) that were gained in the ancestor of all pathogens (Fig. 5a and Supplementary Table 3 and Table 4). Figs. 5b,c,d show the evolutionary histories of the three orthogroups. The tree topology supports a common ancestry of the homologous genes at the base of the *Liberibacter* pathogens and their origins were likely from bacteria other than the non-pathogenic *Liberibacter* ancestor.

### The gene-loss and gene-gain events might contribute to the establishment of endophytic habitats of pathogens

To understand the functional significance of these two types of orthogroups, we performed Gene Ontology and KEGG pathway analysis. This revealed that many of the genes that were lost in the endophytic pathogens are involved in synthesis and metabolism of several major types of cellular components, including amino acids, phosphate-containing compounds, nitrogen components, and nucleotide acids (Supplementary Figure 1, 2, and Table 5). Strikingly, the whole biosynthetic pathways for Tryptophan and Histidine were missing in the pathogens (Fig. 6). The majority of the biosynthetic genes are clustered together as different operons on the ancestral *L. cresens* genome; however, a complete deletion of several related operons and other genes highlights the dependency of these intracellular pathogens on receiving these nutrients from their hosts.

**Fig. 6.**
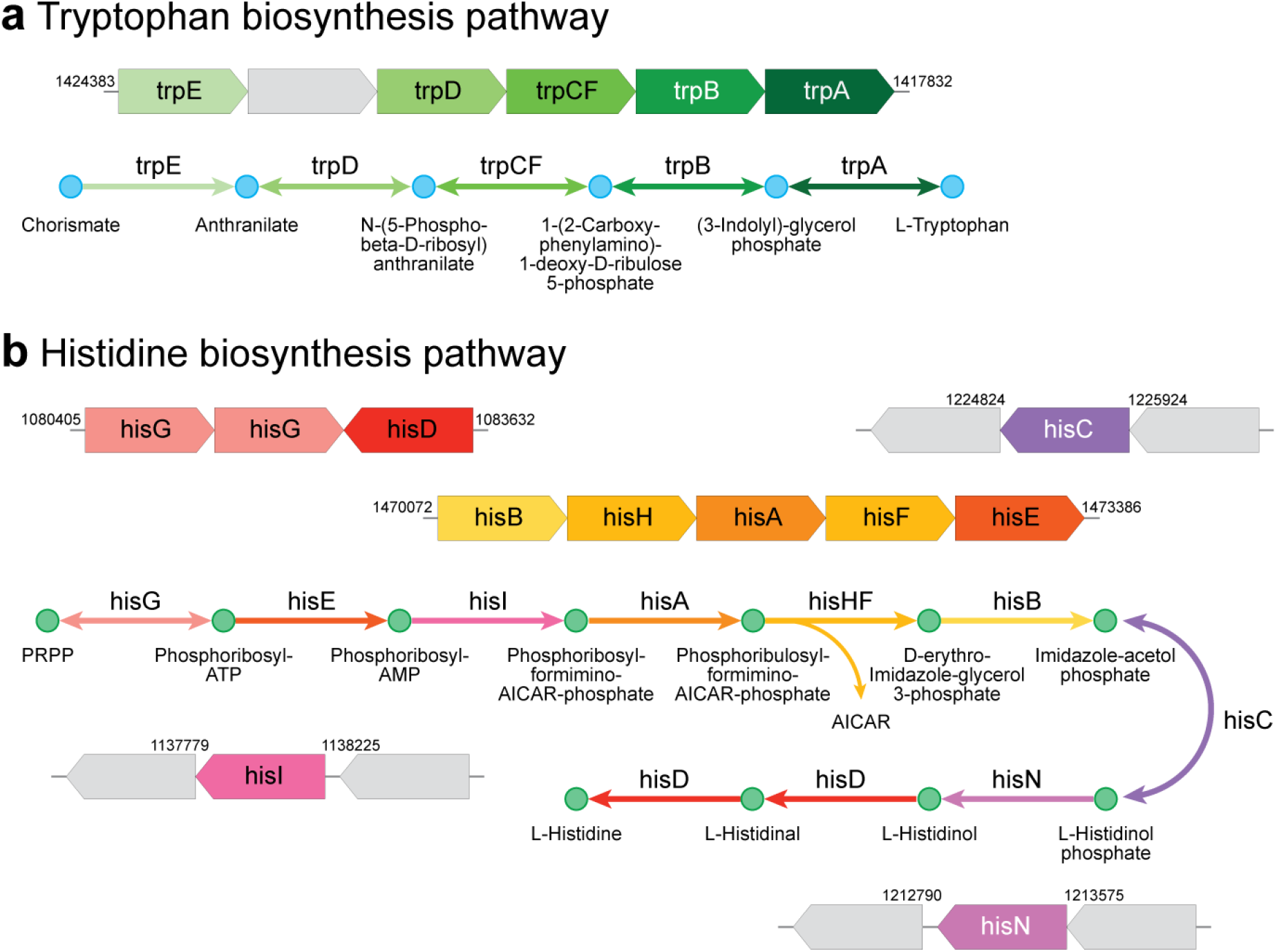
KEGG pathway analysis reveals a complete deletion of a chain of enzymes of the pathways responsible for biosynthesis of Tryptophan (**a**) and Histidine (**b**) in the pathogenic *Liberibacter* species. In each case, operonic structure of biosynthesis enzymes in the ancestral *L. crescens* BT-1 genome is presented above, with the enzyme genes in colored arrow blocks and gene boundaries indicated by genomic locations; the KEGG pathway is shown below, with reaction steps in arrowed lines, colored according to the operonic blocks. The grey blocks indicate the genes that are not associated with the pathway and not lost in pathogenic *Liberibacter* species.

For the genes that were gained in the pathogens, we found that they encode multiple transporters, enzymes involved in DNA/RNA synthesis, regulation, or small molecule metabolism, and several transmembrane proteins (Supplementary Table 4). Some of them might contribute to the pathogen’s ability to adapt and proliferate in the intracellular environment of host cells. For example, the survival protein SurE, is a metal-dependent nucleotide phosphatase that has been shown to be essential for bacterial pathogenesis and survival in the stationary phase and in harsh conditions ^40^. Further, the multiple transporters might play important roles in exchanging chemical components between the pathogens and the plant host ^41^. Thus, both the gene-loss and gene-gain events that happened in the ancestor of *Liberibacter* pathogens seem to have defined the molecular foundation of their endophytic habitats.

### Extensive sequence and structure analyses identify potential virulence factors, including several polymorphic toxins

Since pathogenicity is an acquired trait for *Liberibacter* bacteria, we hypothesize that such ability is attributed to the unique pathogenicity-related genes, that were gained by the ancestor of these pathogens during evolution (or gene-gain events). Therefore, the potential virulence factors, such as toxins and effectors, should be among the gene-gain list that we identified (Supplementary Table 4). However, of the proteins on our list, almost half of them are hypothetical proteins with no functional annotation. To accurately uncover the function of these proteins, we conducted a series of sequence and structure analyses to examine the sequence/structural features of these proteins, dissect their domain components, establish the distant relationship between these domains with the known Pfam domains, and further synthesize the function of the proteins by combining the domain annotations. This systematic analysis has allowed us to identify three groups of proteins as potential toxins/effectors (Supplementary Table 4).

The first toxin group comprises two hypothetical proteins (e.g. WP_012778667.1 and WP_012778668.1 in *C*.L. asiaticus strain gxpsy). By sequence analysis, we found that these two proteins share a unique domain architecture containing two previously un-recognized domains, a PD domain (designated for the conservation of two residues, Pro and Asp; Supplementary Figure 3) and a Ntox52 domain (Supplementary Figure 4). No function has been associated with these domains. However, by studying the proteins containing either the PD or Ntox52 domains, we found that they are also coupled with other domains that are components of polymorphic toxin systems (PTSs). PTSs are a vast class of toxin systems that we have recently identified in bacteria and archaea ^42–45^. They have served as the primary weaponry for many bacterial pathogens ^46–49^ which were exported through different secretion pathways such as T2SS, T5SS, T6SS and T7SS ^42,43,45,50,51^. Despite the extensive diversity of these polymorphic toxins, we were able to dissect the underlying principles of these proteins, including 1) The toxins typically contain multiple modular architectures with N-terminal secretion-related domains, central repeats or linker domains, pre-toxin domains, and C-terminal toxin domains ^42,43,45^; and 2) The toxins display a tremendous polymorphism in domain composition as they diversify through domain recombination or shuffling ^42,43,45^. Therefore, the association of toxin-related domains is the most prominent feature of toxin proteins ^42,43,45^.

We found that both the PD and Ntox52 domains display a typical association with other toxin-specific domains (Fig. 7a). Specifically, the PD domain is frequently coupled with several long N-terminal regions and a number of C-terminal toxin domains including nucleases (REase-1, AHH, EndoU), peptidases (MPTase, Tox-PL4), deaminases, ADP-Ribosyltransferases (ART-HYD3, ART-VIP2), and Anthrax toxA (Fig. 7a), that were identified in our earlier studies ^42,43,45,52^. The Ntox52 domain, on the other hand, is always located at the C-terminus of toxin proteins, a position that the toxin domain typically occupies, and coupled with other toxin-specific central modules, such as RHS and PseudoCD2, in addition to the PD domain. These lines of evidence strongly suggest that the two identified proteins represent novel polymorphic toxins where the PD domain is a pre-toxin module and the Ntox52 is a toxin module. By secondary structure prediction and conservation analysis, the Ntox52 domain is featured by a core structure containing several alpha helices and four beta strands and by several conserved polar residues (Supplementary Figure 4). However, no distant homolog has been found in our profile-based sequence searches.

**Fig. 7.**
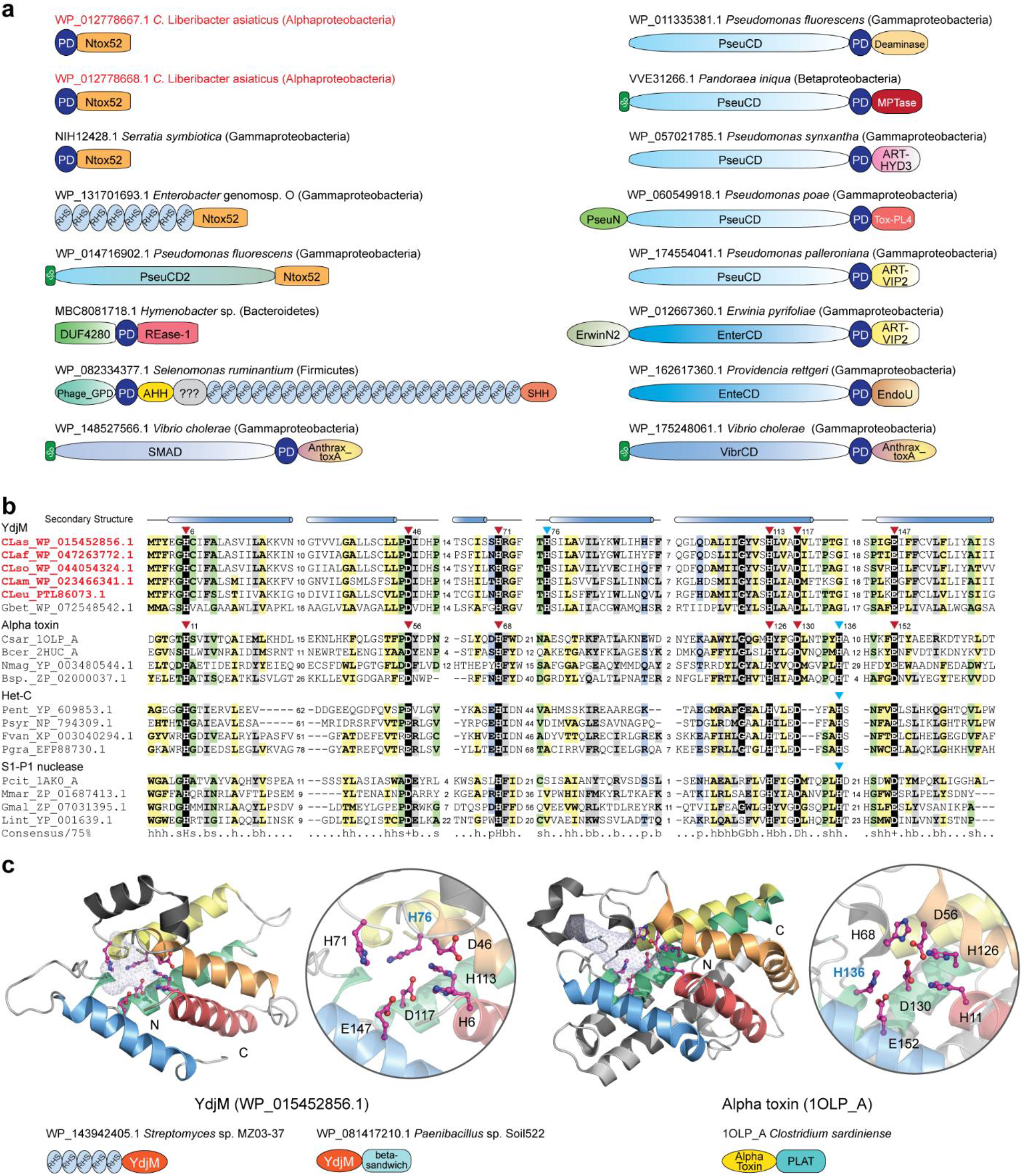
Identification of potential protein toxins. **a** Domain architectures of representative polymorphic toxins containing the PD domain and the Ntox52 domain. The PD toxins of *C*. L. asiaticus were highlighted in red. Domain architectures are labelled by accession numbers, species names and their lineages in bracket. Domain architectures are not drawn to scale. **b** Multiple sequence alignment between the YdjM, alpha toxin, Het-C, and S1-P1 nuclease domains. The conserved catalytic residues are highlighted in black background. **c** Structural comparison of YdjM and the alpha toxin domains. Representative domain architectures of these toxin proteins are shown.

Another potential toxin is the YdjM protein (Fig. 7b). Our profile-profile comparison revealed that it is related to several toxin domains including bacterial alpha-toxin (probability 95% of profile-profile match), a membrane-disrupting phospholipase responsible for gas gangrene and myonecrosis in *C. perfringens* infected tissues ^53^, and the HET-C domain (probability 93% of profile-profile match), that is also a toxin component that we identified in bacterial PTSs and in the fungal nonallelic incompatibility system ^43,54^ (Fig. 7b). Although they share a low sequence identity, these domains display conservation in both the structural composition and the catalytic core (Fig. 7b and 7c). The catalytic core of the known alpha toxin and Het-C domains involves seven conserved residues, six of which are preserved in the YdjM domain. However, our structural modeling suggested that the YdjM-specific His76 serves an equivalent role as His136 of the alpha toxin (Fig. 7c). Further, by analyzing the domain architectures of many other proteins containing the YdjM domain, we found: 1) several bacterial RHS-type toxins use YdjM as their C-terminal toxin module (e.g. WP_143945134) (Fig. 7c); 2) the YdjM domain is predominantly coupled with a novel beta-sandwich domain (Fig. 7c; Supplementary Figure 5), like the alpha toxin, which has a beta-sandwich domain to facilitate toxin localization at the membrane ^53^. Thus, based on these lines of evidence, we proposed that YdjM is a novel phospholipase toxin which might disrupt the membrane of host cells.

The third toxin group includes several proteins belonging to the EEP (Endonuclease/Exonuclease/Phosphatase) family (Fig. 5b and Supplementary Table 5). They contain an N-terminal signal peptide, indicating that they are secreted and therefore might target host cells. The EEP family comprises many enzymes with different activities, from DNA I-like nucleases, endonucleases of retrotransposons, inositol polyphosphatases to phosphodiesterases (Pfam ID: PF03372). By a phylogenetic analysis of the *Liberibacter* EEP related sequences and other known EEP enzymes (Supplementary Figure 7), we found that the *Liberibacter* EEP and related sequences form a distinct clade which is more closely related to a clade of DNA I-like nucleases, which also include *Haemophilus ducreyi* cytolethal distending toxins (1SR4_B)^55^ and *Salmonella* typhoid toxin (4K6L_F)^56^. Therefore, it is possible that the secreted *Liberibacter* EEP proteins act as nuclease toxins targeting the host DNAs.

## Discussion

HLB is the most destructive disease of citrus worldwide and associated with several species of *Candidatus* Liberibacter, a psyllid-transmitted, phloem-limited, alpha proteobacteria. Despite intense scrutiny, molecular mechanisms of the HLB pathogenesis remain to be elucidated. The fact that these pathogens are obligate intracellular bacteria and are not culturable *in vitro* has made experimental, functional studies extremely challenging. Thus, computational mining of available genome information has become a promising strategy to dig into the potential pathogenesis mechanisms of HLB and to guide the directed experimental studies. Here we present an in-depth comparative genomic analysis of several *Liberibacter* species, including those pathogens associated with citrus HLB and potato ZC, along with the ancestral, non-pathogenic *Liberibacter* species, *L. crescens*. We successfully identified the major genomic changes that might contribute to the establishment of endophytic habitats and pathogenicity of the *Liberibacter* pathogens.

One of the major genome variations found by our comparative genomic analysis resides at the prophage loci (Fig. 2 and Fig. 3). Prophages in *Liberibacter* have been extensively studied in the past, but the results are controversial. On the one hand, the primary HLB pathogen, *C*.L. asiaticus, is found to carry two prophage loci whose lytic cycle was activated during bacterial infection in plant ^36^, indicating that the prophage might be involved in pathogenicity or infection. On the other hand, the absence of the prophage in a Japanese strain of *C*.L. asiaticus Ishi-1, which also induced severe symptoms in citrus, suggests that the prophage might be dispensable for pathogenicity. Further, many different prophage loci are identified in both *Liberibacter* pathogens and the free-living ancestor ^30,37^, making it more difficult to understand their functional contribution. To resolve these puzzles, we conducted a unique, comprehensive operonic association analysis to recover all the prophage loci in the *Liberibacter* genomes and systematically annotate the conserved phage components. According to both the prophage composition and phylogeny of the conserved terminase, we were able to classify these diverse prophage loci into four major types, namely LC1, LC2, SC, and UT. This result has clarified several confusions about the presence of the prophage loci in the *Liberibacter* species. For example, our phylogenetic analysis has linked the recently classified type 4 prophages from the *Liberibacter* pathogens ^37^ to the earlier LC2 prophage from the ancestral *L. crescens*^30^. We also uncovered a previously un-detected prophage type, the UT phage, that is present in all the *C*. L. asiaticus strains. Further, we were able to identify the LC2 and UT prophages in the Japanese strain which was previously thought to contain no prophage loci ^5^. Thus, our detailed annotation and classification of prophages has provided a genomic resource for studying the prophages in the *Liberibacter* bacteria.

Importantly, our analysis revealed that these different types of prophages originated through multiple independent gene transfer events to *Liberibacter* at different evolutionary stages (Fig. 4c). However, it is only the SC-type prophage that was acquired by the ancestor of the pathogens. Thus, this evolutionary association suggests that the acquisition of the SC phage might contribute to the emergence of pathogenicity in these pathogens. Indeed, many toxins of bacterial pathogens are known to be carried by prophage loci ^33–35^. On average, a typical SC-prophage locus contains about 35 genes. While the majority of these genes encode structural components and enzymes involved in phage replication and transcription, others remain uncharacterized. Therefore, a systematic analysis of the SC-prophage components will be necessary to understand its role in the *Liberibacter* pathogens. To be noted, we observed an unusual divergence between the SC prophage loci from different *Liberibacter* pathogens and different strains of the same pathogen, distinct from other genomic regions and other types of prophages. This rapid evolutionary pattern of the SC prophage suggests that some of its components may have acted at the interface of host-pathogen interactions and gained the upper hand during the evolutionary arms race ^57,58^.

Our comparative genomic analysis also identified genomic (non-prophage) genes that were lost or gained at the ancestor of the *Liberibacter* pathogens. To be noted, of the 323 orthogroups (335 genes) that were lost in all the pathogens, 56 genes were also identified as the culture-essential genes for *L. crescens* in a Tn5 transposon mutagenesis screening experiment ^59^. Functional and pathway analysis revealed that biosynthetic pathways of several essential amino acids in the pathogens were disrupted by gene loss, consistent with the earlier studies ^30,60^. Some of the genes that were gained in the ancestor of the pathogens encode transporters or transmembrane proteins, which may also contribute to the molecular adaptation of the bacteria to endophytic habitats. Further functional characterization of the genes on the gene-loss and gene-gain lists holds promises for advancing our understanding about the nutrient dependence of endophytic *Liberibacter* bacteria and enabling rational design of new strategies to culture them *in vitro*.

More importantly, our detailed sequence and structural analyses have identified several potential protein toxins. Protein toxins or effectors are the major pathogenicity components of bacteria-caused diseases. However, the toxins that are responsible for HLB and other related diseases have remained enigmatic for years. Early genome annotation did not reveal any bacterial pathogenesis-related genes in the genome ^5^. Although several studies have attempted to examine the genes encoding small, secreted proteins, candidates thus far are only limited to certain *C*.L. asiaticus strains ^14,16,18–21^. We had also tried to identify toxins in *Liberibacter* by utilizing our previously established toxin domain profiles of polymorphic toxin systems; unfortunately, no candidate was identified. This suggests that the *Liberibacter* pathogens might utilize some novel toxins whose toxin domains are not yet identified. Indeed, by studying the hypothetical proteins that were found to follow a gene gain pattern, we were able to uncover three new toxin groups. Importantly, the newly identified toxin domains are also fused with other typical domains of the polymorphic toxins. This strongly supported their role as toxins in the *Liberibacter* pathogens. In addition to the above toxin candidates, other hypothetical proteins in the gene-gain list may also be potential toxins. One such example is the protein (WP_015452959.1) that was recently found to cause cell death in the leaves of *N. benthamiana*^61^.

This research expands our recent efforts in using computational means to dissect the molecular mechanisms of complex organismal interactions ^42,43,45,62^. Organismal or species interactions, either inter-specific, intra-specific, or those between pathogen and host, are the mainstay of Life ^43^. Driven by the evolutionary arms race, the proteins mediating these interactions, such as toxins, effectors, or virulence factors, typically evolve rapidly and are difficult to identify using traditional experimental and computational methods ^63^. The major contribution of our research in this regard is to use dedicated protein domain-centric analysis strategies, comparative genomics, and evolutionary theory to identify such toxin/effector components and to formulate the principle of the systems behinds these interactions ^42,43,62,63^. Previously, this approach has led to the discovery of several distinct classes of conflict systems, including bacterial polymorphic toxin systems involved in kin selection and bacteria-host interactions ^42,43,45^, Crinkler-RHS (CR) effector systems at the interface of eukaryotic pathogen/symbiont-host interactions ^62^, nucleotide-centric conflict systems ^64^, DNA modification systems deployed in phage-bacteria interactions ^65^, and viral pathogenicity factors involved in coronavirus-host interactions ^57,58^. Many of our computational predictions in protein function and the organizational principles of the systems have facilitated the creation of new concepts and directions in several research fields and have been later experimentally validated, including the recent discoveries of the enzymatic function of animal and plant TIR proteins ^66,67^, adenosine methylation enzymes ^68,69^, type III CRISPR-Cas systems ^70,71^, and novel immunoglobulin proteins and ion channel proteins in SARS-CoV-2 ^57,58^, in addition to many work on PTSs ^42,43,45^.

As an extension to understand the interactions between bacterial pathogens and their host, we present our identification of the elusive pathogenicity factors in the *Liberibacter* bacteria. This knowledge will open doors to developing and deploying new strategies for the detection of *Liberibacter* pathogens and treatment of the associated diseases. Both the SC prophage loci and the genes gained in the ancestor of the pathogenic strains, especially the newly identified toxin groups, can serve as biomarkers for specific detection of the pathogens. The association of these genes with pathogenic species suggests that some of them may play a role in pathogenesis. This hypothesis can be tested directly using targeted, functional experiments. We envision that future research informed and enabled by our genomic analysis holds great promises to elucidate the pathogenesis mechanisms of *Liberibacter* bacteria, leading to long-sought solutions for curing HLB and other related diseases.

## Methods and material

### Genomes that were investigated in this study

A total of 12 genomes of *Liberibacter* species were collected from the nucleotide database of the National Center of Biotechnology Information (NCBI). Their genome annotations including the coding sequences (CDS) and RNA genes were extracted from NCBI GenBank files. The encoded protein sequences were retrieved from the NCBI protein database. The detailed information of the genomes used in this study can be found in Supplementary Table 1.

### Pair-wise genome comparisons

To systemically identify the genomic features and variations among *Liberibacter* species, we performed pair-wise whole genome comparisons by using the local TBLASTX ^72^ program with the cut-off e-value of 0.001 serving as the significant threshold. The results were visualized using Artemis Comparison Tool (ACT) ^73^ with a cut-off matching sequence length at 500 bp, and modified using Adobe Illustrator. The *C*.L. europaeus genome has 15 contigs and in order to compare it to other genomes, we linked these contigs based on the order of their accession numbers: PSQJ01000001.1, PSQJ01000002.1, PSQJ01000004.1, PSQJ01000005.1, PSQJ01000006.1, PSQJ01000007.1, PSQJ01000008.1, PSQJ01000009.1, PSQJ01000010.1, PSQJ01000011.1, PSQJ01000012.1, PSQJ01000013.1, PSQJ01000014.1, PSQJ01000015.1 and PSQJ01000003.1.

### Protein sequence search and analysis

To collect protein homologs, iterative sequence profile searches were conducted using the Position-Specific Iterated BLAST (PSI-BLAST) program ^72^ against the non-redundant (nr) protein database of NCBI with a cut-off e-value of 0.005 serving as the significance threshold. Similarity-based clustering was performed by BLASTCLUST, a BLAST score-based single-linkage clustering method (ftp://ftp.ncbi.nih.gov/blast/documents/blastclust.html). Multiple sequence alignments (MSA) were built by the KALIGN ^74^, MUSCLE ^75^ or PROMALS3D ^76^ programs, followed by careful manual adjustments based on the profile–profile alignment and the secondary structure information generated by the JPRED program ^77^. A consensus method was used to calculate the conservation pattern of the MSA based on different categories of amino acid physicochemical properties developed by Taylor in 1986 ^78^. Consensus is calculated by examining each column of the MSA to determine whether a threshold fraction (either 75% or 80%) of the amino acids belongs to a defined category. Then the MSA was colored using the CHROMA program ^79^ based on the calculated consensus sequence and further modified using Adobe Illustrator or Microsoft Word. The HHsearch program was used for profile-profile comparison ^80^. Signal peptide and transmembrane region prediction was detected using the Phobius program ^81^. Potential open reading frames were detected using the ORFfinder program (https://www.ncbi.nlm.nih.gov/orffinder/).

### Molecular phylogenetic analysis

We used both the Maximum Likelihood (ML) analysis, implicated in the MEGA7 programs ^82^, and Bayesian Inference (BI), implemented in the BEAST 1.8.4 program ^83^, to reconstruct the phylogenetic relationships of genes and proteins.

To infer the phylogeny of *Liberibacter* species, we selected three genes, including 16S rRNA, 23S rRNA, and DNA polymerase A, collected their homologs from *Liberibacter* bacteria and other related sequences using BLASTN or BLASTP programs, generated MSAs which were further analyzed using the ML analysis. A GTR model was applied to the RNA sequences and a JTT model was applied to the protein sequences. Initial tree(s) for the heuristic search were obtained automatically by applying Neighbor-Join and BioNJ algorithms to a matrix of pairwise distances estimated using the Maximum Composite Likelihood (MCL) approach, and then selecting the topology with a superior log-likelihood value. For both data types, a discrete Gamma distribution was used to model evolutionary rate differences among sites (four categories). A bootstrap analysis with 100 repetitions was performed to assess the significance of the phylogenetic grouping.

To reconstruct the evolutionary history of terminases, we first applied the ML analysis in which a WAG model with a discrete gamma distribution (four categories) was used to model rate heterogeneity among sites. A bootstrap analysis with 100 repetitions was performed to assess the significance of the phylogenetic grouping. We also applied the BI analysis with a JTT model and a discrete Gamma distribution (four categories). Markov chain Monte Carlo (MCMC) duplicate runs of 10 million states each, sampling every 10,000 steps was computed. Logs of MCMC runs were examined using Tracer 1.7.1 program ^83^. Burn-ins were set to be 2% of iterations.

All trees with the highest log-likelihood from the ML analysis were visualized using the MEGA7 program ^82^. The tree is drawn to scale, with branch lengths measured in the number of substitutions per site. The bootstrapping values from ML analysis and/or the Posterior values from Bayesian inference analysis are shown next to the branches.

### Gene neighborhood analysis of the prophage loci

In order to identify the prophage loci, we utilized the gene neighborhood analysis. Unlike many earlier studies ^25,30,37^, which relayed on the existing phage database and sequences, our analysis is based on the operonic association to comply with the extreme diversity of prophage loci. As the terminase is the core component of prophage, we used it as a marker to identify potential prophage loci on the genome. We collected the upstream and downstream gene neighbors of the terminase from the NCBI GenBank files. All protein sequences were clustered using the similarity-based BLASTCLUST program (ftp://ftp.ncbi.nih.gov/blast/documents/blastclust.html). Protein clusters were further annotated with the conserved domains which are identified by the hmmscan program searching against Pfam ^80,84^ and our own curated domain profiles.

### Identification of the genes gained or lost at the ancestor of pathogenic species

To identify the genes that were gained or lost at the ancestor of all *Liberibacter* pathogens, we first utilized the OrthoFinder v2.2.3 program ^39,85^ with default parameters to infer groups of orthologous gene clusters (orthogroups) among all 12 *Liberibacter* proteomes based on protein homology detection by Diamond ^86^ and MCL clustering ^87^. Orthogroup comprises genes of related species that evolved from a common gene ancestor by speciation ^88^. Given the common ancestry of all *Liberibacter* species, the majority of the genes should share common ancestors and be preserved in all these species, whereas the genes that were gained or lost at the ancestor of the *Liberibacter* pathogens will display a different presence in either pathogenic *Liberibacter* species or non-pathogenic ancestor. Therefore, we extracted these orthogroups using a custom python script based on the following criteria: 1. Orthogroups with gene loss: the ones present in non-pathogenic *L. crescens* but not in any of the pathogenic *Candidatus* Liberibacter species; 2. Orthogroups with gene gain: the ones that are found in at least 4 *Candidatus* Liberibacter species (a total of 5 species used in this study), but not in *L. crescens* genomes.

Importantly, by conducting case-by-case phylogenetic analyses, we realized that there were still some false positive and false negatives in the ortholog clustering result. To overcome this methodological limitation, we utilized another strategy given the special situation of *Liberibacter* species. For ortholog clustering, the gene transfers between bacterial genomes are the major obstacle. In the case of *Liberibacter* bacteria, all pathogenic species are endophytic bacteria and their life cycle is entirely restricted in the host cells. Therefore, they have little chance to exchange genes with other bacteria and evolution of their genes should have been mainly influenced by sequence diversification and their relationship should be readily computationally tractable using pairwise sequence similarity scores. Based on this evidence, we conduced genome-wide BLASTP searches against the NCBI-NR database using all *C*.L. asiaticus proteins as queries. The results showed a good correlation between sequence similarity scores and the evolutionary distance. Thus, by this simple BLAST search, we were able to identify 1) the proteins in the non-pathogenic ancestor that have no close homologs in pathogenic *Liberibacter* species, and 2) the proteins in pathogenic species that have no close homologs in the ancestor.

To validate the above results on gene gain and gene loss, we carried out extensive bootstrapped maximum likelihood phylogenetic analyses using the MEGA7 program for all the identified orthogroups by including both *Liberibacter* proteins and other related homologs that were identified from NCBI-NR database. These three steps led to an identification of 323 orthogroups (about 335 genes) that are only found in non-pathogenic *L. crescens* and 35 orthogroups that are only present in pathogenic bacteria (Supplementary Table 3 and Table 4). The number of gene-loss and gene-gain genes might differ between species or strains due to genome-specific duplications.

### Protein function analysis

We used three levels of analyses to conduct the functional annotation of the proteins that were identified in this study. First, we retrieved the annotation information from the NCBI RefSeq database and Gene Ontology (GO) terms using the Blast2GO program ^89^. Their involvement in the molecular synthesis pathways was determined by mapping the KEGG database ^90^. Second, for those proteins that have no functional annotation, we annotated them using the conserved domains which were identified using the hmmscan program searching against Pfam ^80,84^ and our own curated domain profiles. Finally, for proteins with potential uncharacterized domains, we conducted detailed case-by-case analysis of protein sequences and structures, such as homologous sequence searches (PSI-BLAST) ^72^, multiple sequence alignment analysis (KALIGN, MUSCLE, PROMALS3D)^74–76^, secondary structure prediction (JPRED) ^77^, and sequence-profile/profile-profile searches (HHsearch) ^80^, to dissect their domain architectures, identify the conserved sequence/structural features, and to predict aspects of their biochemical and biological function.

### Protein structure prediction and analysis

The MODELLER (version 9v1) program ^91^ was utilized for homology modeling of the tertiary structure of the YdjM protein by using the alpha toxin (1OLP_A) as a template. The sequence identity between the template and the targets is 9%. Since in these low sequence-identity cases, sequence alignment is the most important factor affecting the quality of model ^92^, the alignment used in this analysis has been carefully built and cross-validated based on the information from HHsearch and edited manually using the secondary structure information. Structural analysis and comparison were conducted using the molecular visualization program PyMOL ^93^.

## Data availability

Authors can confirm that all relevant data are included in the paper and/or its Supplementary Information files.

## Acknowledgements

Y.T., C.W., T.S., H.L., and D.Z. are supported by the Saint Louis University Start-up Fund. T.-F.H. is supported by the National Institute of Food and Agriculture Hatch Project 02413. X.W. is supported by a USDA National Institute of Food and Agriculture Hatch project 1018100, National Science Foundation EPSCoR RII Track-4 Research Fellowship (NSF OIA 1928770), an Alabama Agriculture Experiment Station ARES Agriculture Research Enhancement, Exploration and Development (AgR-SEED) award, as well as a generous laboratory start-up fund from Auburn University College of Veterinary. X.L. is supported by the National Institute of Food and Agriculture Hatch Project 02634.

## Author contributions

D.Z. conceived the overall project. Y.T., C.W., T.S., H.L. performed the phylogenetic analysis. Y.T and D.Z. performed genome comparisons, GO and pathway analysis, protein domain identification, domain architecture analysis, and structure analysis/modeling. Y.T. and C.W. conducted the operonic analysis of prophages and studied the function of phage components. Y.T., C.W., and R.F.D were involved in ortholog analysis. Y.T. and R.F.D. contributed to the data organization. Y.T., X.T., T.F.H., X.W., X.L. and D.Z. contributed to the design of the analysis strategies. Y.T., H.L., X.T., T.F.H., X.W., X.L., and D.Z. interpreted the results. Y.T. and D.Z. wrote the manuscript. All authors have read, revised, and commented on the manuscript.

## Additional information

Supplementary Information accompanies this paper are available at Nature Communications online.

